# MutSpot: detection of non-coding mutation hotspots in cancer genomes

**DOI:** 10.1101/740944

**Authors:** Yu Amanda Guo, Mei Mei Chang, Anders Jacobsen Skanderup

**Author notes:** These authors contributed equally to this work (Yu Amanda Guo, Mei Mei Chang). Correspondence should be addressed to Anders Jacobsen Skanderup or Yu Amanda Guo.

## Abstract

**Summary:** Recurrence and clustering of somatic mutations (hotspots) in cancer genomes may indicate positive selection and involvement in tumorigenesis. MutSpot performs genome-wide inference of mutation hotspots in non-coding and regulatory DNA of cancer genomes. MutSpot performs feature selection across hundreds of epigenetic and sequence features followed by estimation of position and patient-specific background somatic mutation probabilities. MutSpot is user-friendly, works on a standard workstation, and scales to thousands of cancer genomes.

**Availability and implementation:** MutSpot is implemented as an R package and is available at https://github.com/skandlab/MutSpot/

**Supplementary information:** Supplementary data are available at https://github.com/skandlab/MutSpot/

## Introduction

Cancer is a genetic disease arising from (driver) mutations that give cancer cells a selective advantage to proliferate and invade. Early cancer genomics studies have mainly focused on the protein-coding regions of the genome. However, even with thousands of cancer exomes sequenced in the past decade, identification of putative driver mutations in the coding regions has still not saturated in many cancer types (Chang, et al., 2016; Lawrence, et al., 2014). Importantly, mutations in the non-coding DNA that constitutes the other 98% of the human genome is even less explored. Tumor whole genome sequencing is however gaining popularity and a recent study of over 2,500 tumor whole genomes by the ICGC/TCGA Pan-Cancer Analysis of Whole Genomes Network (PCAWG) estimated that up to 25% of all tumors harbor non-coding driver mutations (Sabarinathan, et al., 2017). There is therefore a pressing need to develop statistical methods that can leverage these large datasets to predict driver mutations in the non-coding DNA.

Current tools designed to identify non-coding drivers are based on mutation recurrence within regulatory elements (Juul, et al., 2017; Lochovsky, et al., 2015), predicted functional impact of somatic mutations (Mularoni, et al., 2016), or a combination of these approaches (Dhingra, et al., 2017; Hornshoj, et al., 2018). However, existing methods are designed to explore mutations within defined regulatory regions, such as promoters, enhancers or UTRs, therefore ignoring the rest of the non-coding genome. As such, a typical non-coding cancer driver detection method evaluates less than 5% of the 3 million bases sequenced in a WGS experiment for signs of positive selection. Furthermore, by restricting the analysis to annotated regulatory regions, current tools will miss non-coding drivers that create *de novo* regulatory elements in regions of unannotated DNA. For example, non-coding mutation hotspots upstream of *TAL1* and *LMO1* in T-cell acute lymphoblastic leukemia lead to the formation of *de novo* MYB binding sites that drives the over-expression of *TAL1* and *LMO1* oncogenes (Hu, et al., 2017; Mansour, et al., 2014). Here, we present MutSpot, an R package that systematically and unbiasedly scans the entire genome for mutation hotspots with statistical evidence of positive selection.

## Methods and implementation

To accurately detect mutation hotspots, MutSpot builds a background mutation model that corrects for known covariates of mutation probability, such as local nucleotide context, replication timing, and epigenomic features (Guo, et al., 2018) (**Fig. 1a**). Separate background mutation models are built for single nucleotide variants (SNVs) and small insertions and deletions (indels), as they arise from different mutational processes. Local nucleotide context features are automatically computed by MutSpot based on the list of mutations provided by the user. For SNVs, sequence features include the nucleotide type (A/T or C/G), and tri-nucleotide (1bp flanks) and penta-nucleotide (2bp flanks) contexts of each mutation. For indels, sequence features include the presence of poly-A, poly-T, poly-G and poly-C sequences at the indel sites. MutSpot uses 135 default epigenetic features including transcription factor binding profiles from ENCODE (Zerbino, et al., 2015), the mean replication timing profile of 13 ENCODE cell lines(Hansen, et al., 2010), and predicted APOBEC editing sites that could drive passenger hotspots (Buisson, et al., 2019). Additional epigenomic features can be provided in the bigwig or bed format. MutSpot also calculates the local mutation rate in 100kb genomic bins to correct for additional unexplained regional variation in mutation rates. The most predictive features of mutation probabilities are selected by a LASSO logistic regression model (see **Supplementary Methods**). To account for inter-patient heterogeneity, MutSpot corrects for the mutation burden of individual tumors. Additional patient specific features such as mutation signatures and cancer subtypes can also be integrated into the model. Finally, MutSpot fits a logistic regression model over all positions in the genome to estimate patient- and position-specific background mutation probabilities.

**Figure 1.**
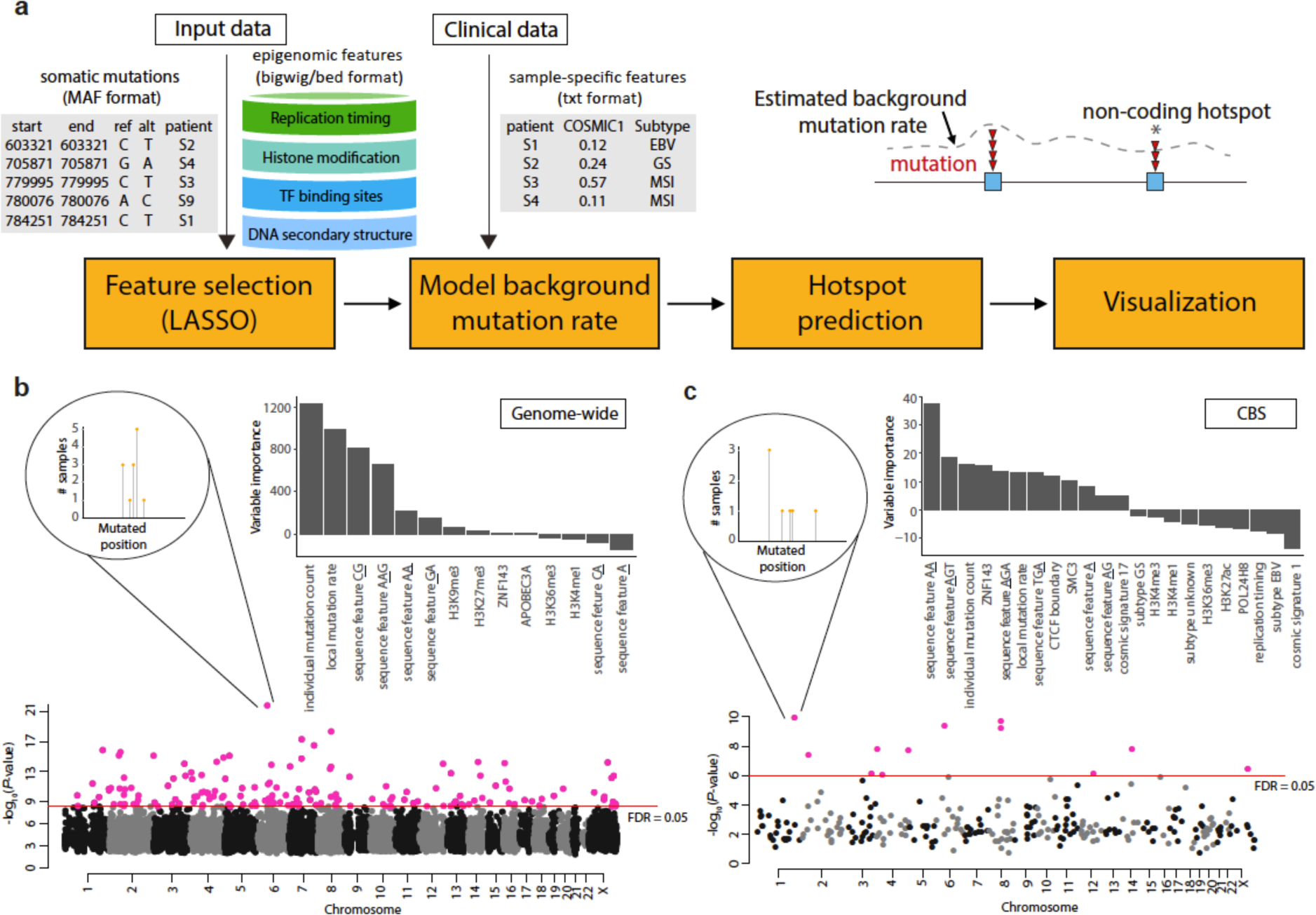
**(a)** MutSpot analysis workflow. **(b-c)** For each analysis, MutSpot outputs 3 types of descriptive figures: a Manhattan plot of the hotspots identified, a feature importance plot of the background model and lollipop plots of the top hotspots. Figures produced by MutSpot from **(b)** a genome-wide analysis and **(c)** a CBS specific analysis on 168 gastric cancer tumors. Hotspots with fdr<0.05 are labeled in magenta.

To identify mutation hotspots, MutSpot evaluates the mutation recurrence for *l*-bp regions genome-wide with at least *n* mutations (default *l=*21, *n*=2). The p-value of mutation recurrence is computed using a Poisson binomial model that accounts for varying mutation rates across different patient tumors (Guo, et al., 2018; Melton, et al., 2015). As mutation hotspot detection can be sensitive to recurrent artifacts, MutSpot excludes problematic regions, such as poorly mappable regions and immunoglobin loci, from the analysis.

## Application examples

MutSpot can be used to detect mutation hotpots either genome-wide or in user defined regions. In the genome-wide discovery mode, MutSpot fits a genomic background model and scans for mutation hotspots across the whole genome. In the regional discovery mode, MutSpot fits a background model specific to the user defined regions, e.g. promoters, and predicts hotspots in the specified regions only. While the genome-wide mode provides a comprehensive scan of the entire genome, the regional mode can be advantageous when the mutational processes in the regions of interest are very different from the genomic background. To demonstrate the utility of the regional analysis, we ran MutSpot on 168 non-hypermutated gastric cancer whole genomes to detect SNV hotspots 1) genome-wide and 2) in regions comprising CTCF binding sites (CBS)(Guo, et al., 2018). In each analysis, MutSpot outputs a Manhattan plot of the detected hotspots and a barplot of the Z-values (quantifying association with mutation rate) of the selected features in the fitted background model (**Fig. 1b and c**). CBSs are known to be hypermutated in gastrointestinal cancers, with a distinct mutation spectrum enriched in A>G and A>C substitutions (Guo, et al., 2018; Katainen, et al., 2015). In the genomic background mutation model, CpG dinucleotides, individual tumor mutation counts and local mutation rate are among the top predictors of mutation probability. On the other hand, consistent with the current knowledge, MutSpot identifies AA dinucleotide as the most important predictor of mutation probability in the CBS-specific model. 24 mutation hotspots at CBSs are identified in the genome-wide model, however, only 8 remains significant in the CBS-specific model that corrects for the elevated background mutation rate and unique mutation spectrum at CBSs.

As it can be computationally expensive to fit genome-wide models with multiple covariates, sparse matrices were implemented to minimize memory usage and a multi-threading option is available to reduce the compute time. MutSpot takes less than 3 hours on a 4 core machine for genome-wide hotspot discovery in 200 tumors, and it can be scaled up to process thousands of tumors on a standard work station (**Supplementary Fig. 1**).

## Discussion

While most existing non-coding driver discovery tools consider sets of predefined features in the background mutation model, MutSpot offers the flexibility to incorporate any genomic or clinical covariate into the background model. This allows the user to include tissue specific epigenetic features for the cancer type of interest, and newly discovered mutational biases into the background mutation model. As our current knowledge of the mutational processes and biases is far from complete, new insights in the processes underlying somatic mutations will further improve the accuracy of hotspot detection.

In conclusion, MutSpot is a user friendly tool for end-to-end non-coding mutation hotspot identification from cancer genomes. As increasing number of cancer whole genomes becomes available, MutSpot can facilitate the discovery of novel drivers in the non-coding genome to further our understanding of tumor biology.

## Acknowledgements

This work was supported by Singapore National Medical Research Council grant OFIRG15nov072. We would like to thank Probhonjon Baruah, Danliang Ho, Weitai Huang, Zhong Wee Poh, and Julie Solacroup for discussion during the development and testing of MutSpot.

